# The *Amidella* clade in Europe (*Amanita* Pers., *Basidiomycota*: *Amanitaceae*): clarification of the contentious *Amanita valens* and the importance of taxon-specific PCR primers for identification

**DOI:** 10.1101/2022.02.15.479696

**Authors:** Ricardo Arraiano-Castilho, Ana Cristina Silva, Carlos Vila-Viçosa, Mário Rui Castro, Luís Neves Morgado, Paulo Oliveira

**Affiliations:** Department of Ecology and Evolution, University of Lausanne, Biophore building, 1015 Lausanne, Switzerland; Instituto Nacional de Investigação Agrária e Veterinária, I.P., 2780-159 Oeiras, Portugal; BIOPOLIS Program in Genomics, Biodiversity and Land Planning, CIBIO, Campus de Vairão, Portugal; MHNC-UP - Museum of Natural History and Science of the University of Porto - PO Herbarium; Praça Gomes Teixeira; 4099-002 Porto; Portugal CIBIO (Research Center in Biodiversity and Genetic Resources) - InBIO (Research Network in Biodiversity and Evolutionary Biology), University of Porto; Campus Agrário de Vairão; Rua Padre Armando Quintas; 4485-661 Vairão; Portugal; Biology Department, Faculty of Sciences, University of Porto, Rua do Campo Alegre, s/n, 4169-007 Porto, Portugal; Convevo Portugal Av. António Augusto de Aguiar, N.19 4. Direito - Sala B 1050-012 Lisboa, Portugal; Metyis Management Consulting Alpha Tower | 7th & 8th floor De Entree 69 1101 BH Amsterdam The Netherlands; MED – Mediterranean Institute for Agriculture, Environment and Development. Mitra College, Apartado 94, 7006-554 Évora, Portugal; CIBIO (Research Center in Biodiversity and Genetic Resources) - InBIO (Research Network in Biodiversity and Evolutionary Biology), University of Porto; Campus Agrário de Vairão; Rua Padre Armando Quintas; 4485-661 Vairão; Portugal

**Keywords:** *Amanita*, *Amidella*, *Amanita pseudovalens*, *Amanita ponderosa*, *Amanita curtipes*, *Amanita lepiotoides*, taxonomic probes, evolutionary convergence

## Abstract

The species in genus *Amanita* section *Amidella* form a well-defined clade, but some taxa remain difficult to discriminate. In particular, the concept of *Amanita valens* (E.-J. Gilbert) Bertault remains controversial. To understand the phylogenetic placement of a set of collections from South Portugal with a novel nrDNA barcode, we have obtained nrDNA sequences for previously unassessed type collections. The taxon formerly described as *Amanita curtipes* f. *pseudovalens* Neville & Poumarat is interpreted as a separate species, *Amanita pseudovalens* (Neville & Poumarat) Arraiano-Castilho *et al*. comb. et stat. nov., and is genetically indistinct from the Portuguese collections, thus clarifying their taxonomic context. However, our collections are morphologically and ecologically distinct (respectively, ellipsoid to oblong basidiospores and association with *Cistus* on acidic schist soils), and are proposed as a new variety, *Amanita pseudovalens* var. *tartessiana* Arraiano-Castilho *et al*.. These developments also enable a better diagnosis of *Amidella* taxa in Europe, a progress that is most decisive for the late Winter to Spring season collections, and identification keys are proposed. However, the co-occurrence and morphological similarity of the new variety, in comparison with the prized edible *Amanita ponderosa* Malençon & Heim, could leave some collections of either taxa unresolved. Thus, a molecular marker approach was developed, to provide a clear and cost-effective identification aid to complement the keys. The proposed diagnostic tools can be applied toward a review of European *Amidella* taxa chorology from existing records, conserved materials, and future collections. Evolutionary convergence may contribute to the determination difficulties in the *Amidella* clade.

## Introduction

*Amidella* E.-J. Gilbert was originally proposed as a genus in the *Amanitaceae*, to encompass a group of species with basidiomes initially white, some of them unchanging and others transitioning to pinkish, ochre or brown tones with maturation, or where rubbed, presenting a non-striated pileus, very friable partial veil leaving a fugacious annulus on the stipe and appendiculate remains in the pileus margin, a thick membranous saccate volva with a friable inner layer, and ellipsoid to subcylindical amyloid spores (Neville & Poumarat 2004). Gilbert soon made it a subgenus of *Amanita* Pers., and currently the name applies, as a section within subgenus *Lepidella*, only to the species that change colour, while the unchanging ones (*Amanita ovoidea* (Bull.) Link and allies) belong now to section *Roanokenses* (Cui et al. 2018, Riccioni et al. 2019). Worldwide there are close to 30 *Amidella* species in several continents, and the type species is the North American *Amanita volvata* (Peck) Lloyd.

In Neville & Poumarat’s (2004) review of European *Amidella*, three species are recognised: *Amanita curtipes* E.-J. Gilbert, *A. lepiotoides* Barla, and *A. ponderosa* Malençon & R. Heim. For each of these species, they created divergent forms: on one hand, *A. lepiotoides A.* f. *subcylindrospora* Neville & Poumarat, for the high variability of spore morphologies in this species; on the other hand, *A. ponderosa* f. *valens* (E.-J. Gilbert) Neville & Poumarat and *A. curtipes* f. *pseudovalens* Neville & Poumarat, because of the forms of intermediate size between the small *A. curtipes* and the typically robust *A. ponderosa*.

The epithet *valens* was originally created by Gilbert as a variety of *A. lepiotoides*, before *A. curtipes* and *A. ponderosa* were known, and later raised to the species rank by Bertault, to fill the size gap between these two; however, the spore morphology of *A. valens* (E.-J. Gilbert) Bertault differs so much from the original form described by Gilbert, that Neville & Poumarat (2004) proposed that the epithet *valens* was being applied to two taxa, the original as the form of *A. ponderosa* mentioned above, in line with the opinions of Malençon (Neville & Poumarat 2009) and Pinho-Almeida (1994), while Bertault’s *A. valens* was reclassified as *A. curtipes* f. *pseudovalens*. A consensus on this matter ought to be met with genetic information on the type specimens, thus the present study investigates the nrDNA from the three divergent forms created by Neville & Poumarat, supporting the concepts of *A. ponderosa* f. *valens* and (provisionally) of *A. lepiotoides* f. *subcylindrospora*, but demonstrating that *A. curtipes* f. *pseudovalens* is a well separated species, forming a new combination, *Amanita pseudovalens* (Neville & Poumarat) Arraiano-Castilho *et al*. comb. et stat. nov..

Furthermore, we describe the Portuguese collections belonging to *A. pseudovalens* as a new variety, *Amanita pseudovalens* var. *tartessiana* Arraiano-Castilho *et al*. var. nov., and due to its very difficult separation from smaller specimens of the prized edible *A. ponderosa*, a molecular marker approach was developed, to provide robust determinations.

## Materials and methods

### Collections

Field sampling was conducted in Southern Portugal during two spring seasons (2010 and 2015) and complemented with herbarium collections, including type specimens from taxa that remained unsequenced (Fig. 1, Table 1). The Spring 2010 collections are from the Barrancos (36.16 N, 7.01 W; 10 samples) and Portel (38.27 N, 7.65 W; 10 samples) counties. The 2015 collections comprise 12 samples from Luzianes (Odemira county) and one from Monte Carvalho (Portalegre county). The Luzianes sites were designated A (37.6072 N, 8.4497 W) and B (37.6033 N, 8.4469 W), and C for a neighbouring site (precise location unknown). The single Monte Carvalho collection was made on a path, with a sandy texture on the surface, at the edge of a *Quercus suber* L. grove located in Alto da Quinta Nova (39.3361 N, 7.4158 W). Additional materials: three *Amanita ponderosa* specimens offered by traditional collectors in Spring 2015, from uncharacterised locations (Ode13–15); and three *Amanita curtipes* collections made during Spring 2010 and Autumn 2016 in the Mitra university campus, Évora (38.53 N, 8.02 W), in association with evergreen oaks on sandy soil. The 16 specimens from the 2015 field campaign were deposited in the PO Herbarium (Natural History and Science Museum of the University of Porto, MHNC-UP), with consecutive codes between PO-F2132 and PO-F2147.

**Fig. 1.**
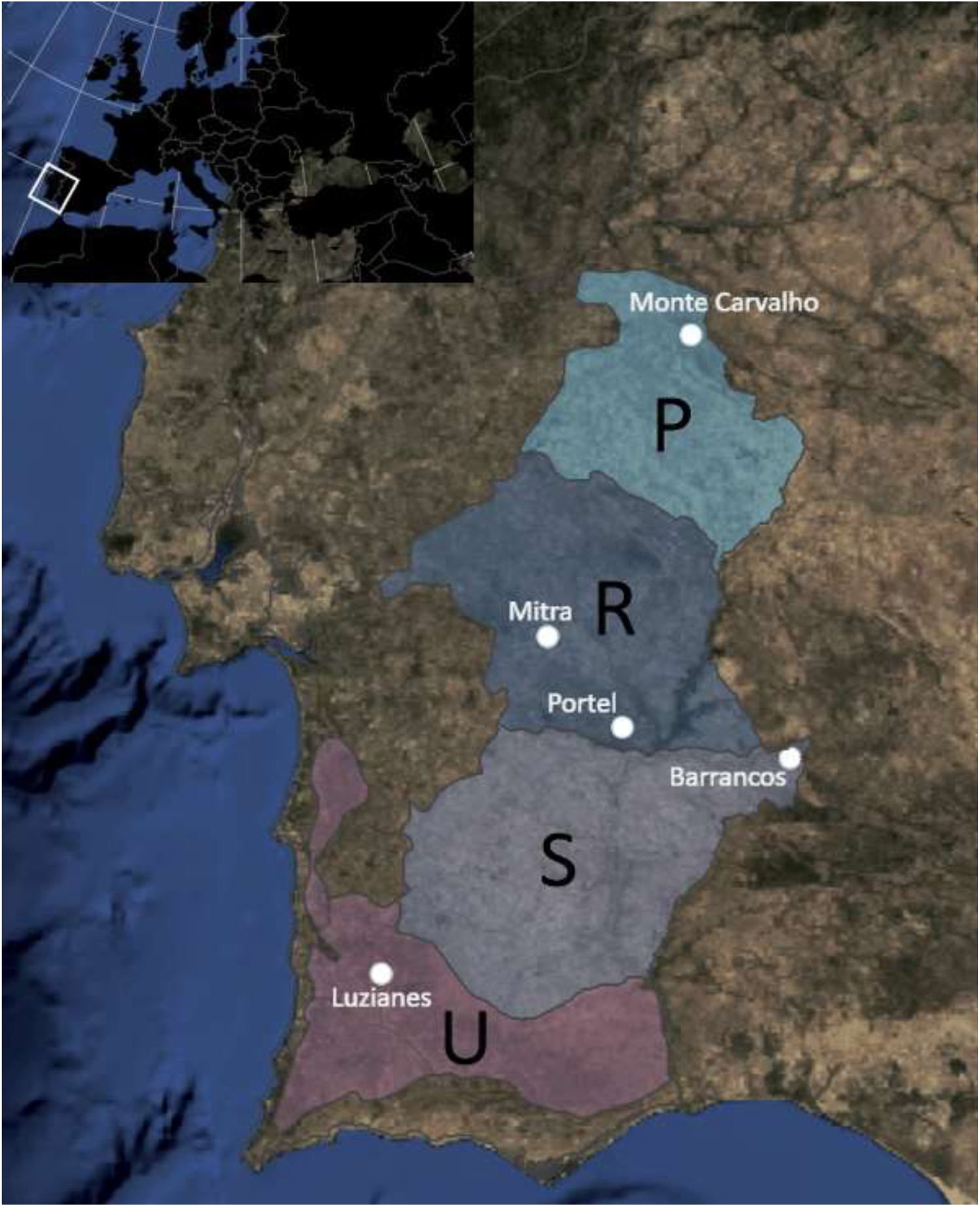
Locations in Continental Portugal for the collections used in this study (Table 1). The letters designate Landscape Units Alto Alentejo (P) Alentejo Central (R), Baixo Alentejo (S) and Serras do Algarve e do Litoral Alentejano (U). Source: DGT, Carta de Unidades de Paisagem (CUP), https://www.dgterritorio.gov.pt/dados-abertos. The area is outlined on the inset with a white rectangle.

**Table 1.**
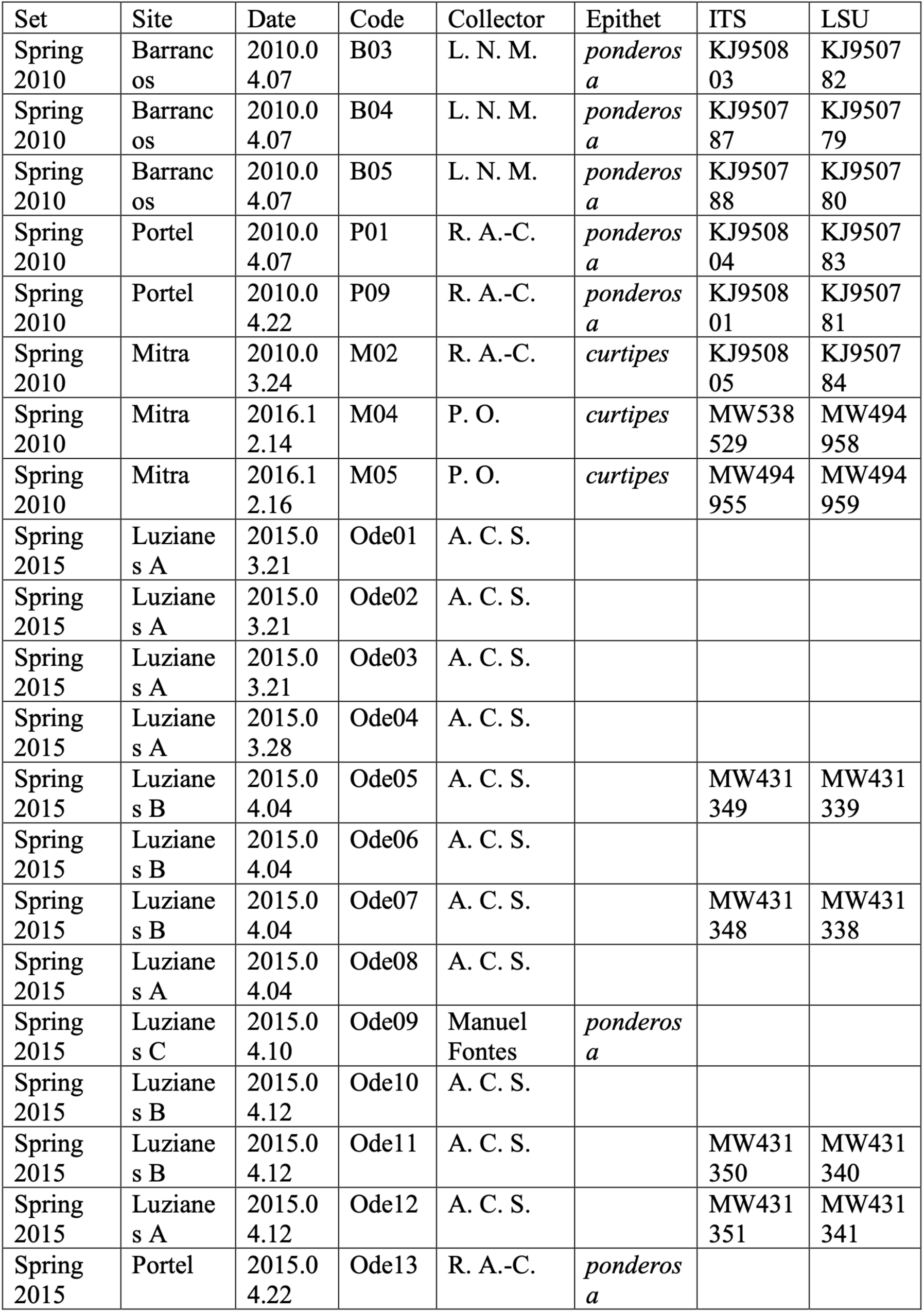
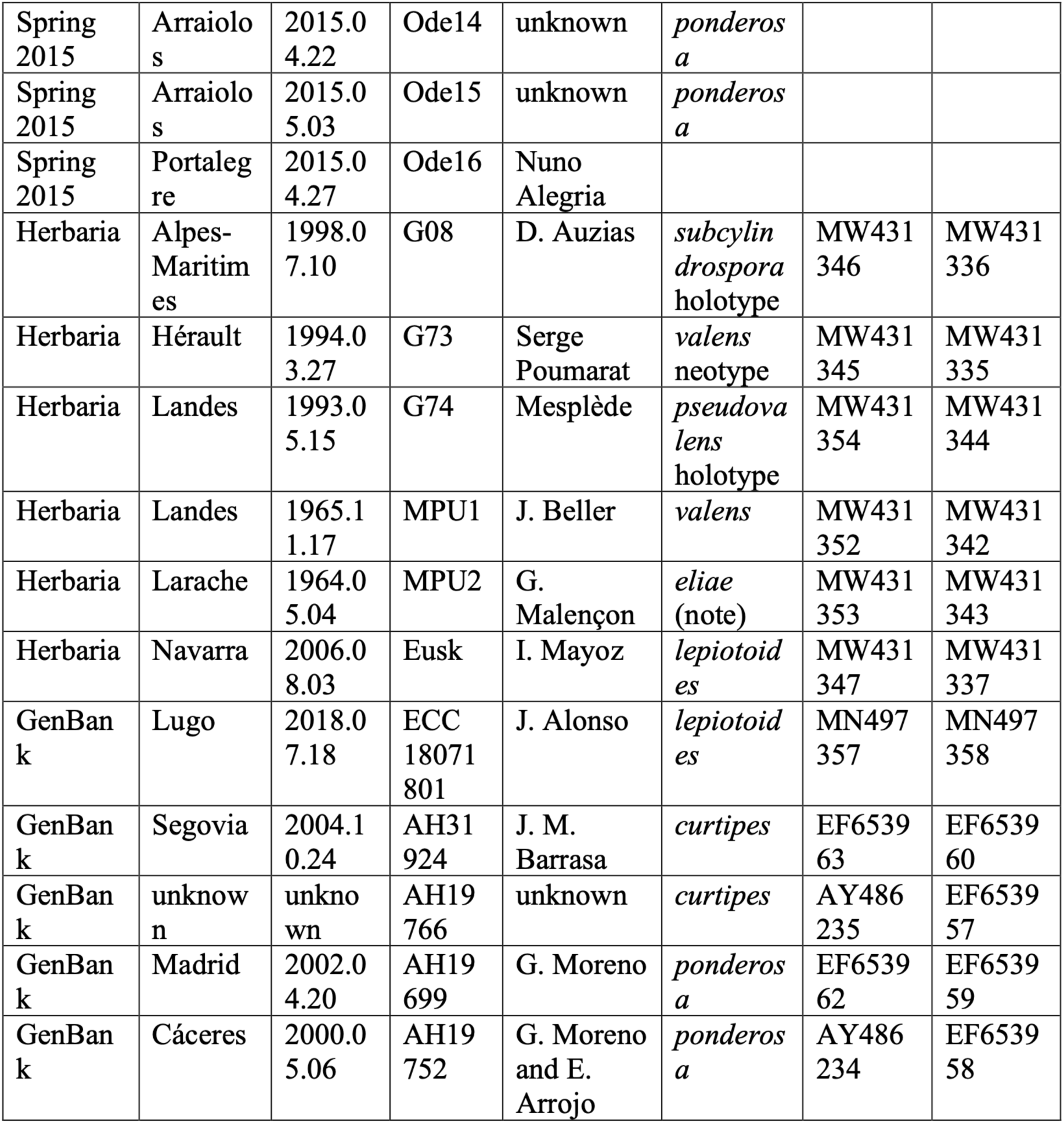
Collections under study, the last two columns the ITS and LSU sequences, as indicated. show the accession numbers for Note: as reported by Neville & Poumarat (2004, page 654), the MPU2 collection was interpreted by Malençon as Amanita curtipes.

The herbaria materials comprised type specimens cited by Neville & Poumarat (2004) ceded by the Conservatoire & Jardin Botaniques de la Ville de Genève (Geneva, Switzerland http://www.ville-ge.ch/musinfo/bd/cjb/chg): the *Amanita lepiotoides* f. *subcylindrospora* holotype (barcode G00126408, SIB: 270556/1), the *Amanita ponderosa* f. *valens* neotype (collection number SGP 94.03.206, barcode G00561773, SIB: 383077/1) and the *Amanita curtipes* f. *pseudovalens* holotype (SGP 93.05.15.203, barcode G00561774, SIB: 382650/1). Materials from two further collections were ceded by the University of Montpellier - Institute of Botany (MPU) herbarium (https://science.mnhn.fr/institution/mnhn/search), one labelled *Amanita valens* from the Landes department, France (catalogue number MPUC00085) and the other labelled as *Amanita eliae* from Larache, Morocco (G. Malençon collection number 5225, catalogue number MPUC00038). Finally, one *Amanita lepiotoides* from the Aranzadi Science Society herbarium (reference ARAN-Fungi 5040268), collected in Navarra and reported by Arrillaga & Mayoz (2005), Spain, was provided by Pedro Arrillaga.

### Characterization of the sites

The Barrancos and Portel sites belong to a geomorphological unit dominated by Paleozoic siliceous materials, mainly schists or granites. The territory has a dry to sub-humid ombrotype, over a thermomediterranean to mesomediterranean thermotype belt (Capelo & Vila-Viçosa, 2020; Costa et al. 1998; Monteiro-Henriques et al. 2016; Rivas-Martínez et al. 2017). The potential vegetation is a oak woodland co-dominated by cork oak (*Quercus suber*) and round-leaf oak (*Q. rotundifolia* Lam.) that, by anthropic action, result in an open landscape and patch-like mosaic known as Montado/Dehesa (Blanco-Castro et al. 2005). All collections were performed in skeletal soils that are disturbed by agricultural and silvicultural activities. The prevailing understory is composed of grasslands and shrubland associations dominated by *Cistaceae*, *Lamiaceae* and *Fabaceae* species (e.g., *Cistus ladanifer* L., *C. salviifolius* L., *C. crispus* L., *Lavandula* Sect. *Stoechas* Ging., *Genista* L. and *Ulex* L.). These shrub associations belong to *Ulici eriocladi-Cistetum ladaniferi* and *Genisto hirsutae-Cistetum ladaniferi* (Costa et al. 2012) and they represent the late regressive and pioneer stages resulting from erosion of the soil upper layer, intensive grazing or fires (Castro & Freitas, 2009; Mendes et al. 2015).

The collections from Luzianes sites A and B sites were on compact, eroded and acidic clay soils derived from schistose bedrock, at the edge of *Eucalyptus globulus* Labill. plantations. The vegetation type is a shrubland dominated by *Cistus ladanifer* (*Cisto ladaniferi – Ulicetum argentei*) with an estimated vegetation cover of 70-80%, representing an early regressive stage of cork oak forests (*Lavandulo viridis – Quercetum suberis*, Quinto-Canas et al. 2010).

### Mycological descriptions

Descriptions of the basidiomes collected during Spring 2015 were made on site and completed in the laboratory by following the guidelines of a specifically designed observation form. Basidiome photographs were taken on site and on arrival to the laboratory. Reagents used for microscopy were prepared as described in Clémençon (2009). Basidiospore measurements were made from spore print samples preferably, or from lamellae preparations. Basidiospores were measured with a 100× objective, and basidia with a 40× objective. All measurements were made using calibrated microscope eyepieces, for the most part an AM-423X Dino-Eye USB digital camera coupled with image processing DinoCapture software (AnMo Electronics Corp., Taiwan). Statistical analyses and testing were made using JASP (JASP Team, 2020; https://jasp-stats.org). Reference sporographs were drawn based on the guidelines of Tulloss (1984), plotting basidiospore lengths and widths, and using the values in Neville & Poumarat (2004). The colour reactions to 10% FeSO_4_ were observed under the dissecting microscope, on rehydrated samples of stipe context.

### DNA extraction, PCR, sequencing and phylogenetics

DNA was extracted from basidiome samples using a modification of the method for filamentous fungi described by Stirling (2003). Amplification of the nuclear ribosomal DNA internal transcribed spacers (ITS) was made using different combinations of primers, primarily the NSA3-NLC2 (Martin & Rygiewicz 2004) or the V9D-LS266 (Gerrits van den Ende & de Hoog 1999) primer pairs. For the initial 1 Kb of nuclear large subunit (26S) ribosomal DNA (LSU), the LR0R-LR5 primer pair (Vilgalys & Hester 1990; Rehner & Samuels 1994; Hopple & Vilgalys 1999) was used. All amplifications yield a single product with at least 1 Kb, using a protocol of 30 cycles with annealing temperatures of 61, 57 and 54 degrees C, respectively, in the presence of 0.8 mM dNTPs and 1.5 mM MgCl_2_. The herbaria materials produced very fragmented (and possibly nicked) DNA. Thus, successful amplifications were obtained mostly with shorter segments or, to avoid the amplification of fungal contaminants in some extracts, with taxon-specific primers (Osmundson et al. 2013; Table S2).

The PCR solutions were extracted with chloroform and ethanol-precipitated in the presence of ammonium acetate, using linear polyacrylamide as carrier (Fregel et al. 2010) and sent to a sequencing service (STAB VIDA, Portugal). The sequencing chromatograms were inspected to trim the sequence ends and analyse ambiguous positions, using Applied Biosystems Sequence Scanner (Thermo Fisher Scientific, USA) and Ugene (Okonechnikov et al. 2012). Contigs were assembled using the CAP3 (Huang & Madan 1999) module in Ugene.

Sequence alignments were made using the MUSCLE (Edgar 2004) module in Ugene, and phylogenetic analyses using MEGA (Kumar et al. 2018), with tree rendering from the Newick export using the iTOL web tool (Letunic & Bork 2019). In all cases, the partial deletion option was selected at minimum 50% site coverage (i.e., allowing the aligned sites with fewer than 50% alignment gaps, missing data, and ambiguous bases). A search for the Species Hypothesis codes at the UNITE database version 08FU (Kõljalg et al. 2013) was made based on the ITS sequences obtained.

### Molecular markers

Taxon-specific PCR primers targeting the ITS and LSU regions were developed for the quick detection of *Amanita pseudovalens* among *A. ponderosa* collections (Table 2 and Table S1). The strategy assumes the following: i) DNA extracted from fresh basidiomes, ii) PCR primers at concentration 0.25 μM each, iii) one of the primers is general for fungi, and the other is taxon-specific, iv) the taxon-specific primer has the lower melting temperature (IDT OligoAnalyzer tool, https://www.idtdna.com/calc/analyzer/) and v) the PCR annealing temperature is equal to the taxon-specific melting temperature. Negative controls use water, and positive controls include one *A. pseudovalens* reference DNA, and/or a parallel reaction to detect false negatives, either by using primers specific for *A. ponderosa* or matching two general fungal primers.

**Table 2.**
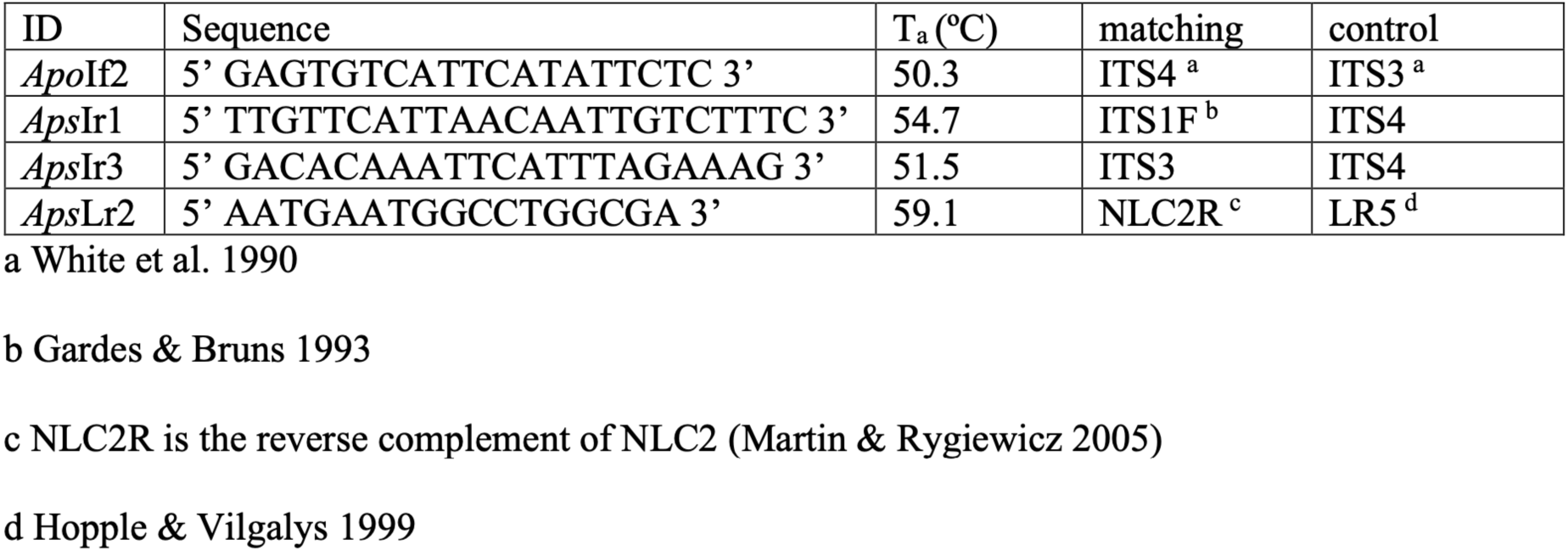
List of selected taxon-specific primers, showing the sequences, annealing temperatures (T_a_), matching and positive control universal primers; *ApoIf2* is for *Amanita ponderosa*, the remainder are for *A. pseudovalens*.

## Abbreviations

nrDNA: nuclear ribosomal DNA region

ITS: nrDNA internal transcribed spacer

LSU: nuclear ribosomal large subunit DNA region

NUTS: Nomenclature of territorial units for statistics

## Results

### Phylogenetic analyses

The phylogenetic analysis of the concatenated ITS-LSU regions revealed four distinct clades corresponding to *A. ponderosa*, *A. lepiotoides*, *A. curtipes* and sister clade of *A. curtipes* containing the type specimen of *A. curtipes* f. *pseudovalens* (Fig. 2). The latter clade also includes the provisionally named *A*. aff. *curtipes* that had been collected in 2010 among other collections of *A. ponderosa* (specimens B03 and P01), along with the MPU and Ode sequences. The same result was obtained separately for the ITS and the LSU regions (not shown). This clade is thus suggested to represent a separate taxon at species level that can be collected both in Southern France and in Portugal, henceforth designated *Amanita pseudovalens* comb. et stat. nov..

**Fig. 2.**
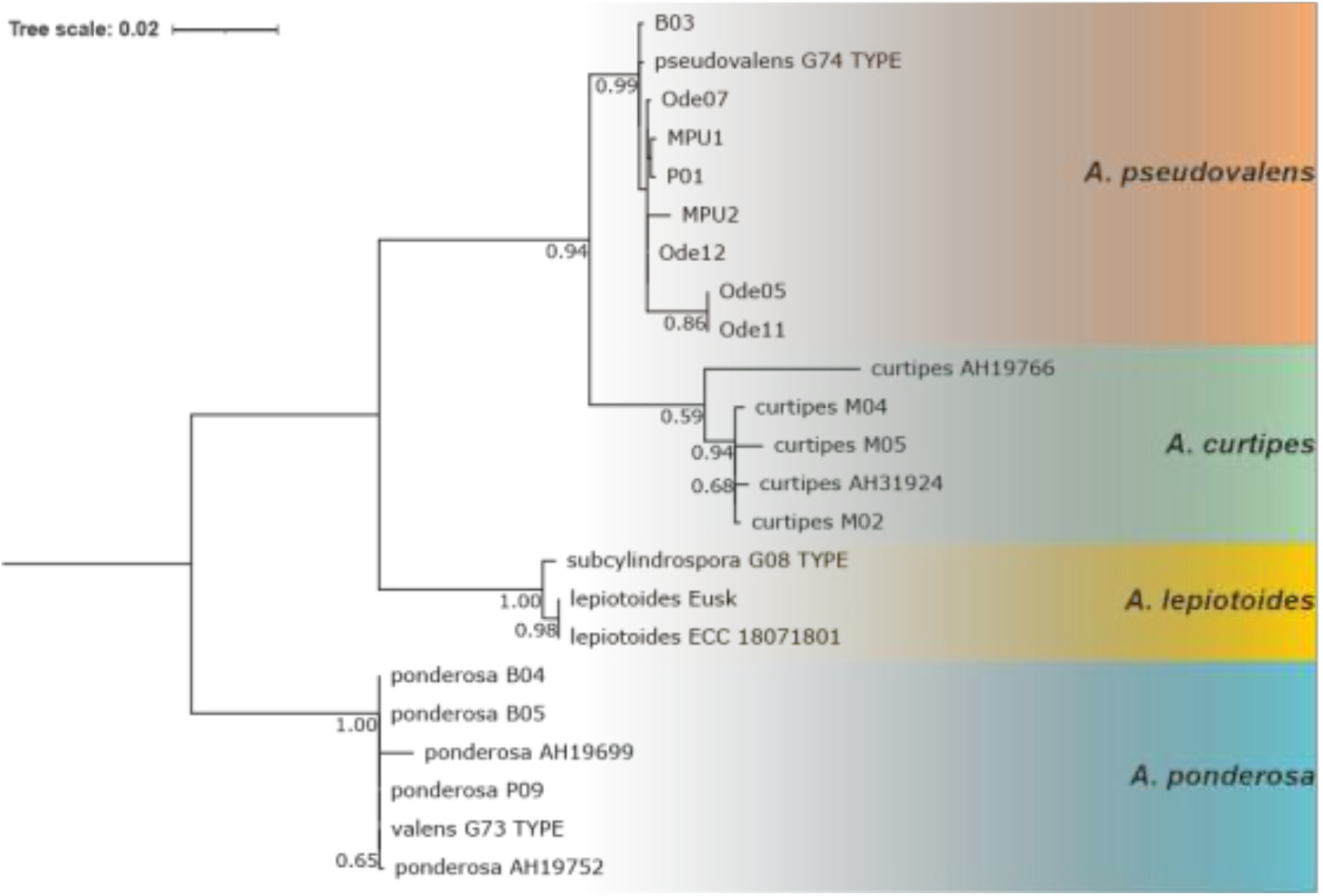
Phylogenetic placement of the type nrDNA sequences, in relation to ITS+LSU sequences from European *Amanita* taxa belonging to the *Amidella* clade (only the terminal epithets are shown). Using a 50% coverage cutoff, a total of 1,546 aligned positions were analysed with the Maximum Likelihood method and the General Time Reversible substitution model (Nei & Kumar 2000), with a discrete Gamma distribution to model evolutionary rate differences among sites (5 categories (gamma parameter 0.2950)), allowing for some sites to be evolutionarily invariable (42.21% sites). The percentage bootstrap support for each node (1,000 replicates) is shown next to the branches. The support for each species clade is 99% (*Amanita pseudovalens* (Neville & Poumarat) Arraiano-Castilho *et al*.), 59% (*Amanita curtipes* E.-J. Gilbert), 100% (*Amanita lepiotoides* Barla) and 100% (*Amanita ponderosa* Maleçon & Heim). The scale indicates 0.02 substitutions per site.

To provide further support to the proposal of a separate taxon at species level, the ITS region was studied for genetic divergence estimates. Using the Net Between Group Distances estimator (Tamura & Nei 1993) in MEGA, the between-node distances for each clade pair were calculated. The net evolutionary divergence (i.e., the number of substitutions per site) between the *curtipes* and *pseudovalens* nodes is positive, albeit lower than other comparisons as expected (Table 3); even without corrections, the divergence estimate (p) for the ITS, between the *curtipes* and the *pseudovalens* nodes, was estimated at 6.4%. The *curtipes* and *pseudovalens* ITS sequences corresponded unambiguously to different Species Hypothesis clusters (Kõljalg et al. 2013) even at the 3% dissimilarity threshold (SH1184304.08FU and SH1184305.08FU, respectively).

**Table 3.**
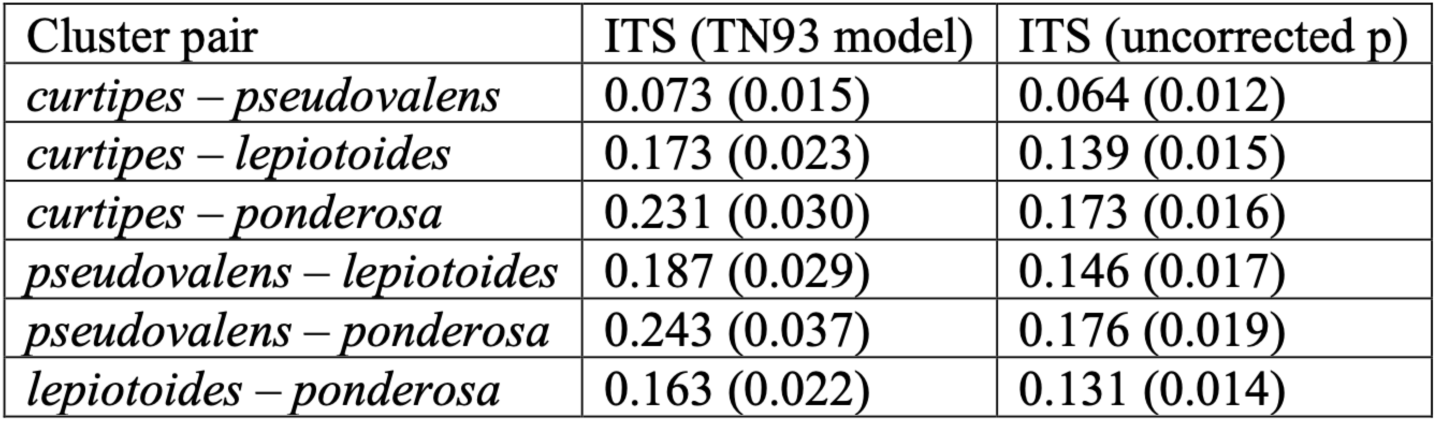
Net evolutionary divergence (base substitutions per site, standard errors in brackets) between groups of sequences (Tamura 1992).

The average Within-Group distance (Tamura-Nei substitution model, Table 4) for the *pseudovalens* clade (containing the Portuguese and French collections) was estimated at 0.0106 nucleotide substitutions per site (± 0.0027 standard error).

**Table 4.** Estimates of average Within-Group distances (nucleotide substitutions per site, standard errors in brackets, Tamura-Nei model) and the Species Hypotheses (SH) for the *curtipes* and the *pseudovalens* clusters defined in this study.

| Cluster | Within-Group | 1.5% SH | 3% SH |
| --- | --- | --- | --- |
| *curtipes* | 0.0031 (0.0013) | SH1562916.08FU | SH1184304.08FU |
| *pseudovalens* | 0.0106 (0.0027) | SH1562918.08FU | SH1184305.08FU |

### Description for the new specimens of *Amanita pseudovalens*

#### Materials included

To describe the new species, we used fresh materials obtained from sites A and B near Luzianes in Spring 2015 (Ode01-12 except Ode09) and the basidiospores from a collection in the Portel county (P01, near Monte Novo, Spring 2010) and another from São Mamede Park near Monte Carvalho, Portalegre county (Ode16, Spring 2015). All specimens used in this description were deposited in the PO herbarium (see Materials and Methods).

### Macroscopy

#### Pileus

Flat to slightly depressed, convex at the margin, expanding to a diameter of 7.5 cm. Most collections whitish *in situ*, turning rose/ochre with either aging, handling, or scratching. A pale grey plaque from the universal veil is frequently present. Some collections present brownish scales close to the margin (Fig 3b-c). Margin thinly appendiculate.

**Fig. 3.**
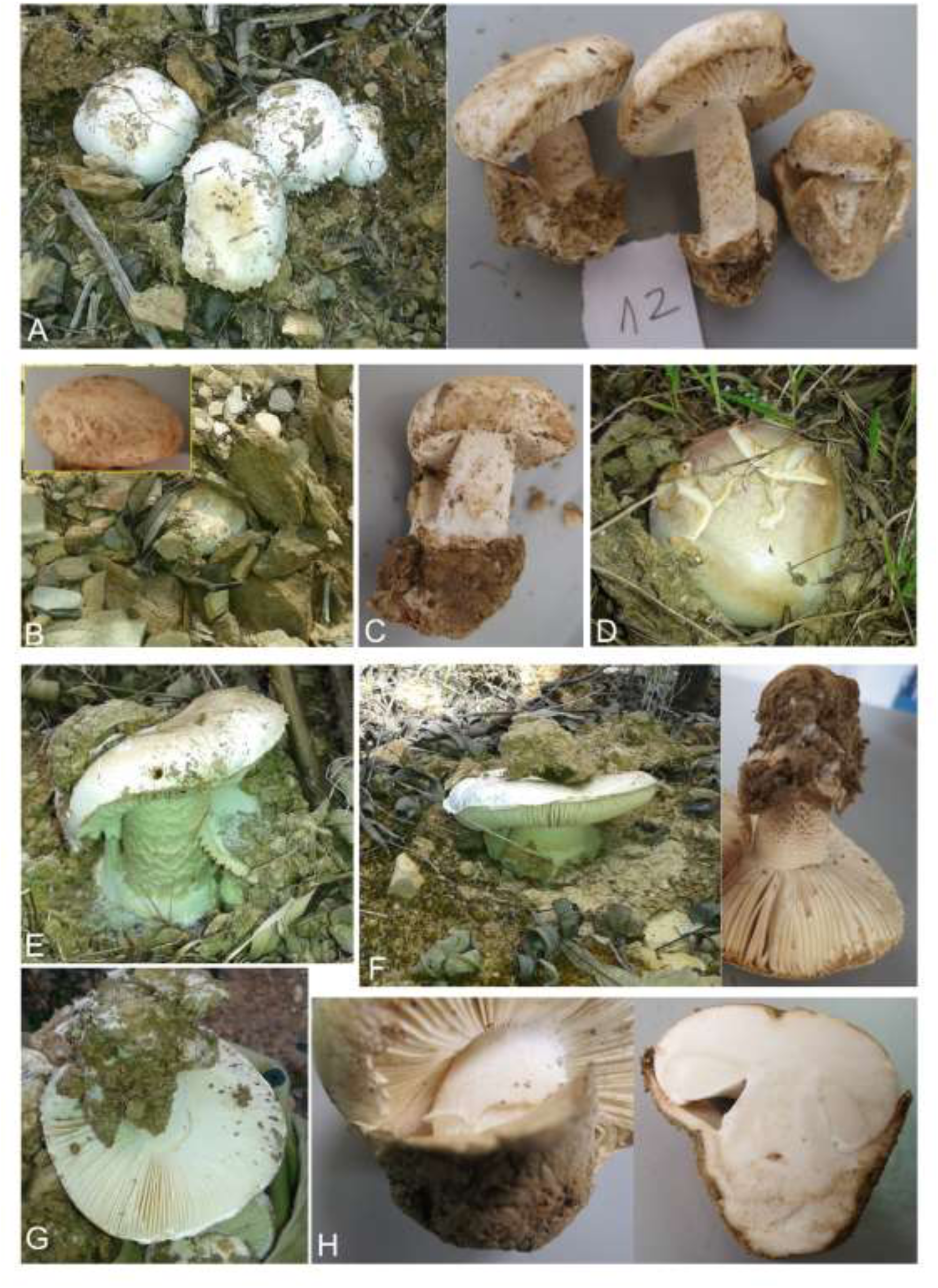
Photographs of *Amanita pseudovalens* specimens from Odemira. Some images taken in the field are matched with corresponding ones taken in the laboratory. **A**, Grouped basidiomes (Ode12) and their appearance upon arrival at the laboratory. **B**, Outcropping away from the vegetation, showing the coarse soil texture in this case (Ode02); inset shows squamulose inner cuticle remains. **C**, Another specimen with cuticle squamules (Ode11). **D**, A specimen with cracked cuticle (Ode05). **E**, An example with stipe squamules (Ode06). **F**, Side view of an emerging basidiome (Ode03) and the same specimen showing the darkened squamules on the stipe. **G**, A view of the hymenophore and the appendiculate pileus margin (Ode02, a specimen different from B). **H**, Details of an immature basidiome (Ode06, a specimen different from E) showing the annulus and insertion of the lamellae (left), and longitudinal section (right).

#### Hymenophore

Adnexed ascending, white, turning rose/ochre with either aging, bruising or scratching, with lamellulae.

#### Stipe

Almost cylindrical, slightly tapering toward the apex, non-bulbous, base obconical. Concolorous with the pileus, with a very fugacious annulus (Fig. 3h). A scale covering can be seen below the annulus region (Fig. 3a/c/e/f). Height not longer than the diameter of the expanded pileus, thickness 2.2 cm at the most.

#### Veil

Universal veil leaving a sac-like thick volva with a lobed margin, pale grey, with an internal ridge raised in contact with the stipe (Fig. 3h); often it remains also as a single pale grey plaque on the pileus. Partial veil leaving a fugacious non-membranous annulus at roughly two-thirds of the stipe height, and narrow remnants on the pileus margin.

#### Context

Concolorous with the surface, homogeneous, relatively compact, non-putrescent. Odour indistinct. Reaction with 10% FeSO_4_ on rehydrated samples from the stipe develops an immediate change to greenish grey that lasts a few minutes.

### Microscopy

#### Basidiospores

White, amyloid, ellipsoid to oblong, average length 11.78 μm, average width 6.97 μm, average length/width ratio (Q) 1.696 (Table 5), overlapping the *A. ponderosa* sporograph but not the one for *A. pseudovalens* (Fig. 4). Due to the lack of spore print, collections Ode01 and Ode16 were not included in the summary calculations. Statistical testing rejected the hypothesis of homogeneity among the collections included in the summary statistics, for all three variables (Fig. S2, Table S3). Indeed, the heterogeneity among collections was the rule (Table 5 and Fig. S3): Ode10 had longer spores and higher Q, bordering on standard *A. pseudovalens* limits, Ode08 had wider spores and lower Q, even more than *A. ponderosa*, while Ode11 had spores of average Q values but small size. Nested ANOVA (within and between sites) also suggested heterogeneity within sites for the three variables, but only for length and width between sites. Site A collections have on average significantly higher values (Table S4).

**Fig. 4.**
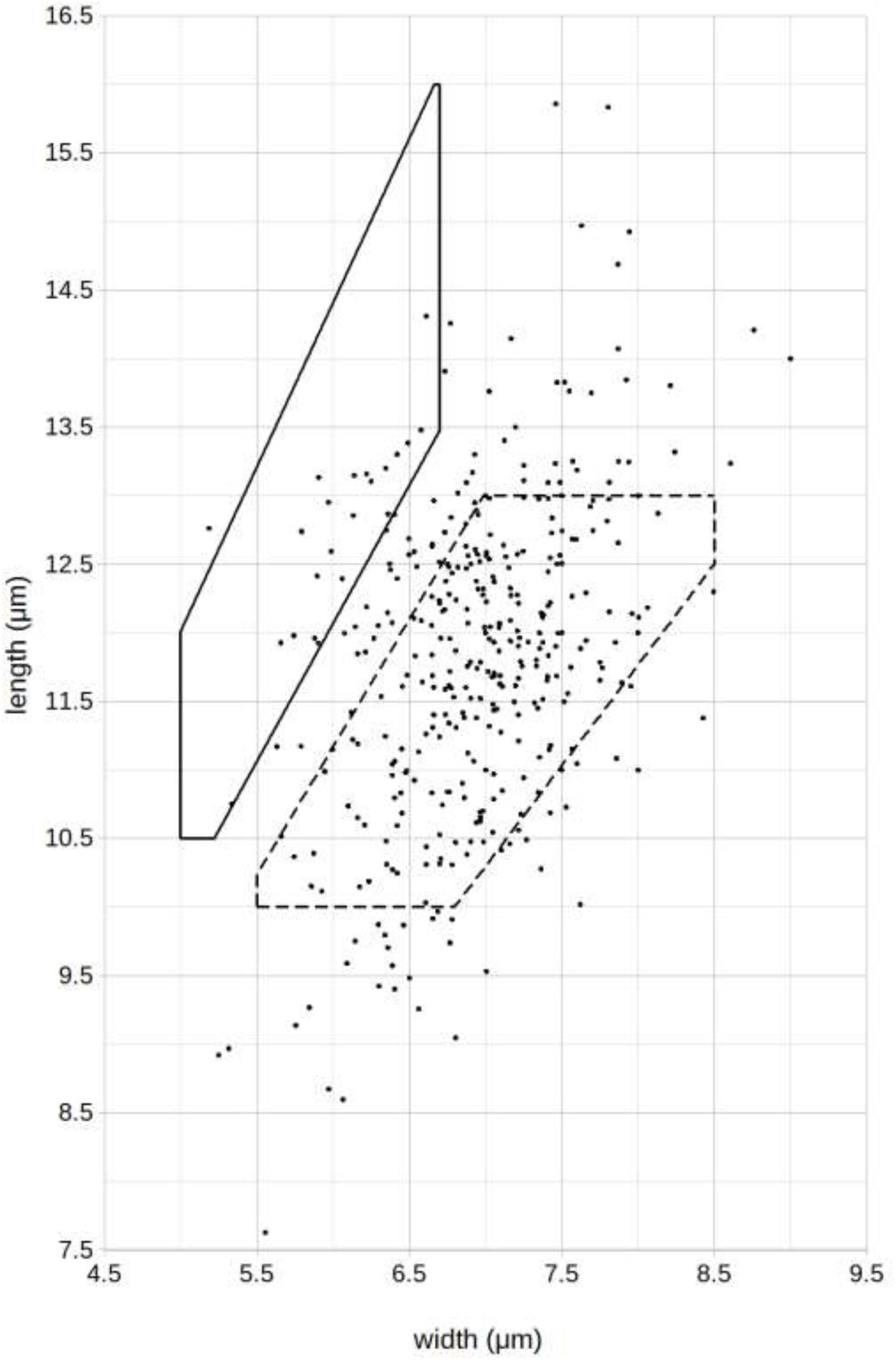
Plotting of the measurements summarised in Table 5 (dots), comparing with the limits of the sporographs for *Amanita pseudovalens* (continuous line) and *Amanita ponderosa* (dashed line).

**Table 5.**
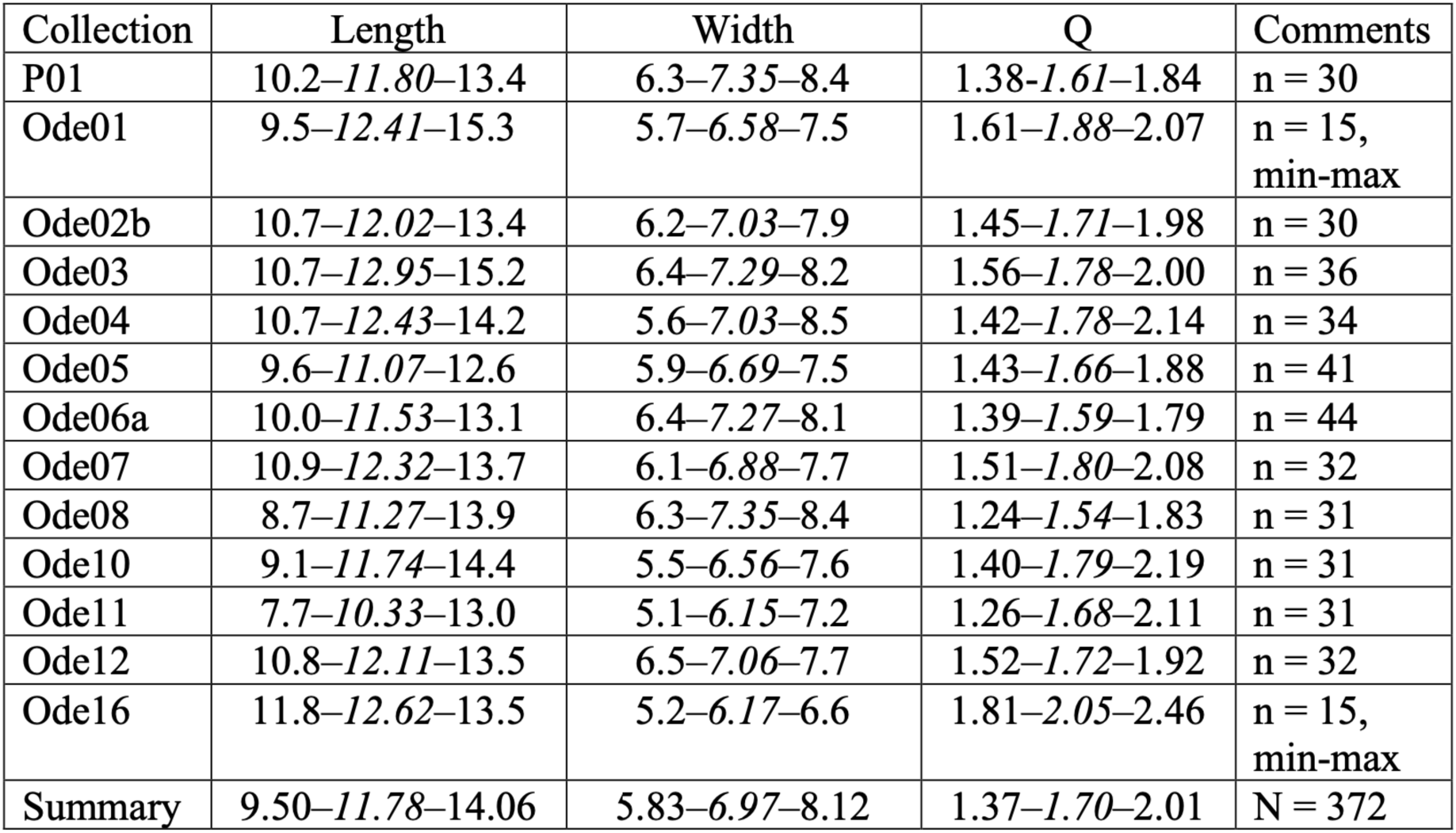
Basidiospore measurements (μm) and Q values, represented as the central 90% distribution intervals (or minimum-maximum ranges (min-max) where indicated), with averages in italic. Summary distributions do not include Ode01 and Ode16 (no spore print available).

| Collection | Length | Width | Q | Comments |
| --- | --- | --- | --- | --- |
| P01 | 10.2–*11.80*–13.4 | 6.3–*7.35*–8.4 | 1.38–*1.61*–1.84 | n = 30 |
| Ode01 | 9.5–*12.41*–15.3 | 5.7–*6.58*–7.5 | 1.61–*1.88*–2.07 | n = 15,<br>min-max |
| Ode02b | 10.7–*12.02*–13.4 | 6.2–*7.03*–7.9 | 1.45–*1.71*–1.98 | n = 30 |
| Ode03 | 10.7–*12.95*–15.2 | 6.4–*7.29*–8.2 | 1.56–*1.78*–2.00 | n = 36 |
| Ode04 | 10.7–*12.43*–14.2 | 5.6–*7.03*–8.5 | 1.42–*1.78*–2.14 | n = 34 |
| Ode05 | 9.6–*11.07*–12.6 | 5.9–*6.69*–7.5 | 1.43–*1.66*–1.88 | n = 41 |
| Ode06a | 10.0–*11.53*–13.1 | 6.4–*7.27*–8.1 | 1.39–*1.59*–1.79 | n = 44 |
| Ode07 | 10.9–*12.32*–13.7 | 6.1–*6.88*–7.7 | 1.51–*1.80*–2.08 | n = 32 |
| Ode08 | 8.7–*11.27*–13.9 | 6.3–*7.35*–8.4 | 1.24–*1.54*–1.83 | n = 31 |
| Ode10 | 9.1–*11.74*–14.4 | 5.5–*6.56*–7.6 | 1.40–*1.79*–2.19 | n = 31 |
| Ode11 | 7.7–*10.33*–13.0 | 5.1–*6.15*–7.2 | 1.26–*1.68*–2.11 | n = 31 |
| Ode12 | 10.8–*12.11*–13.5 | 6.5–*7.06*–7.7 | 1.52–*1.72*–1.92 | n = 32 |
| Ode16 | 11.8–*12.62*–13.5 | 5.2–*6.17*–6.6 | 1.81–*2.05*–2.46 | n = 15,<br>min-max |
| Summary | 9.50–*11.78*–14.06 | 5.83–*6.97*–8.12 | 1.37–*1.70*–2.01 | N = 372 |

#### Basidia

Clavate, with 4 sterigmata, base unclamped, average length 56.9 μm (equal to the median), range 41.0 – 73.4 μm, n = 123. The measurements were obtained from collections Ode02b, Ode05, Ode06a, Ode07, Ode08, Ode10, Ode11 and Ode12, revealing a normal distribution of the global data (Shapiro-Wilk’s W = 0.990, *P* = 0.555). On average, similar basidia sizes were observed across all collections, although Ode11 had a higher average length of 63.9 ± 5.0 μm (Table S5).

### Occurrence

#### Phenology

Late Winter and Spring.

#### Habitat

Mediterranean climate with a dry to sub-humid ombrotype over a thermomediterranean to mesomediterranean thermotype, in association with *Cistus* spp., typically on compact, acidic and eroded soils, corresponding to regressive stages of evergreen oak forests (Fig. 5).

**Fig. 5.**
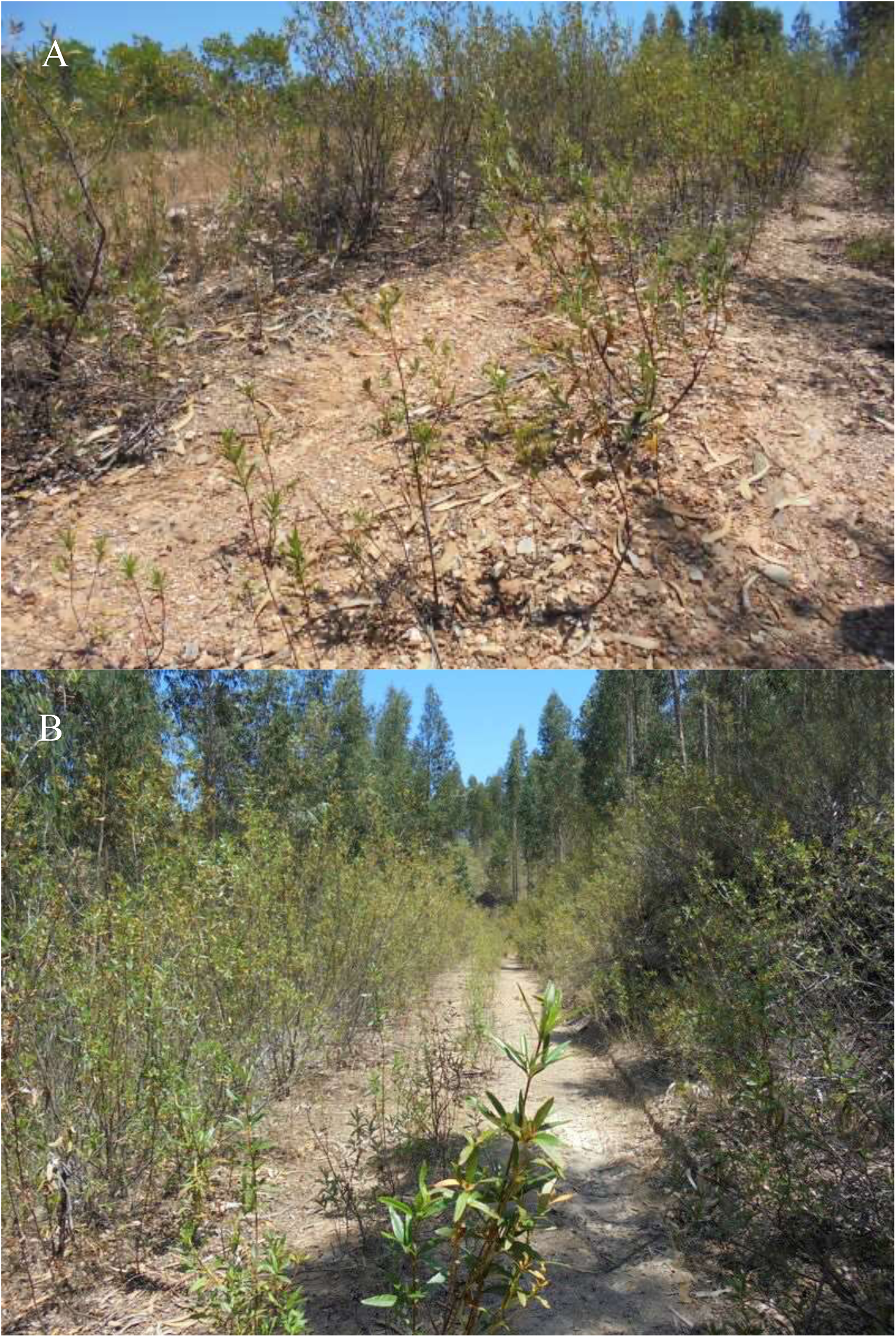
Ground view of the Luzianes locations, showing the *Cistus ladanifer* dominance. **A**, Location A. **B**, Location B. Both photos were taken in Spring 2015 by Ana C. Silva.

#### Distribution

Portugal: reported from the NUTS III regions of Alentejo Litoral, Baixo Alentejo, Alentejo Central, Alto Alentejo.

### Comparison with *Amanita curtipes* f. *pseudovalens* and *Amanita ponderosa*

When comparing the descriptions of all European *Amidella* taxa, based on Neville & Poumarat (2004), as shown in Table 6, the Portuguese specimens of *Amanita pseudovalens* did not conform with the description for the genetically indistinct *Amanita curtipes* f. *pseudovalens* (Neville & Poumarat 2004), diverging in their soil and vegetation affinities and in the microscopy (provisionally, one may also suggest that they tend to be smaller). Therefore, we understand them as representing a separate taxon, however at infraspecific level.

**Table 6.** Comparisons for the described Odemira collections (‘*pseudovalens* Portugal’) with the descriptions made by Neville & Poumarat (2004) for other *Amidella* taxa (‘*pseudovalens* France’ represents *A. curtipes* f. *pseudovalens*).

| Criterion | *curtipes* | *pseudovalens*<br>France | *pseudovalens*<br>Portugal | *ponderosa*<br>f.<br>*ponderosa* | *ponderosa*<br>f. *valens* |
| --- | --- | --- | --- | --- | --- |
| Phenology | Autumn<br>(mostly) | Spring | Winter-Spring | Winter-Spring | Winter-Spring |
| Vegetation | *Pinus*,<br>*Quercus*<br>& others | *Pinus*,<br>*Calluna* | *Cistus* | *Cistus*,<br>*Quercus* | *Cistus* |
| Soil | Sandy | Sandy humic | Schists | Compact | Schists |
| Pileus | 3–7.5 cm | 7–10 cm | 3.5–7.5 cm | 5–15 cm | 4.5–7 cm |
| Stipe | 2–6.7 ×<br>0.5–1.6<br>cm | 6–11 × 1–4<br>cm | 2–6 × 1–2.2<br>cm | 5–13 × 2–<br>4 cm | 8 × 2.5 cm |
| Odour | indistinct | indistinct | indistinct | earthy | recalls<br>young *A.*<br>*ovoidea*<br>("sea",<br>seafood)<br>but<br>weaker |
| Taste | indistinct | initially<br>insignificant,<br>then of<br>hazelnut,<br>slightly bitter | aftertaste<br>slightly bitter | soft | soft |
| Basidiospores | 12.05 ×<br>6.31 µm<br>Q<sub>m</sub> 1.91 | 13.26 × 6.03<br>µm Q<sub>m</sub> 2.20 | 11.78 × 6.97<br>µm Q<sub>m</sub> 1.696 | 11.39 ×<br>6.75 µm<br>Q<sub>m</sub> 1.69 | 10.86 ×<br>7.20 µm<br>Q<sub>m</sub> 1.51 |
| Basidia | 52.1 µm | 52.6 µm | 56.9 µm | 57.6 µm | 59.28 µm |

Moreover, we note that this new taxon is strikingly like *A. ponderosa* in almost every aspect - the obvious difference being the so far unknown occurrence of large and heavy specimens - retaining very little of the affinities with the *A. pseudovalens* collected in France.

### Taxonomic proposals

- *Amanita pseudovalens* (Neville & Poumarat) R. Arraiano-Castilho, A. C. Silva, C. Vila-Viçosa, M. R. Castro, L. Morgado & P. Oliveira var. *pseudovalens* comb. et stat. nov.. Basionym: *Amanita curtipes* f. *pseudovalens* Neville & Poumarat 2004 (Neville & Poumarat 2004: page 656). This taxon corresponds to the French specimens (see Discussion for other possible locations). The basionym diagnosis (Neville & Poumarat 2004) remains unaltered for this variety (Table 6).
- *Amanita pseudovalens* var. *tartessiana* R. Arraiano-Castilho, A. C. Silva, C. Vila-Viçosa, M. R. Castro, L. Morgado & P. Oliveira var. nov.. This taxon corresponds to the Portuguese specimens described in this study. It differs from the autonym by its habitat (with *Cistus* spp., on acidic schist soils), the ellipsoid to oblong, infrequently subcylindric basidiospores, and longer basidia (Table 6). The holotype specimen is Ode12, with Ode02 serving as isotype. A species rank is currently not supported, due to the lack of genetic resolution of the nrDNA sequences. Tartessian refers to an ancient civilization (Тαρτησσός) located in the Iberian South-West (Celestino & López-Ruiz 2016).

Both taxa share, aside from the characters in common with other *Amidella* species, the vernal fruiting season, the medium to small size, and the indistinct odour (Table 6), and can be confirmed with the *Aps* diagnostic PCR primers, as described in Table 2.

### Identification keys

Only the European taxa within the *Amidella* clade are considered. Parts of the keys are from the work of Neville & Poumarat (2004).

1. Pileus remaining convex for most of the development of the basidiome, sometimes with an umbo, diameter notably smaller than the length of the developed stipe, covered at least on the margin with heterogeneous scales from the inner layer of the universal veil, which become brown like the exposed context, stipe base bulbous, margin of the volva leaning towards the stipe, context discoloration moderate to intense. Rare occurrences, widely spaced in time (*Amanita lepiotoides*)…………………………………….2
1. Pileus becoming flat then depressed at the centre, diameter about the same as the stipe length, generally without scales, stipe slightly or not bulbous, margin of the volva free, pale context discoloration, rose then ochre. Occurrence generally annual…………………………………3
2. Context discoloration intense, reddish then dark brown, stipe base distinctly bulbous, basidiospores mostly ellipsoid (Q_m_ < 1.65) ………….*Amanita lepiotoides* Barla f. *lepiotoides*
2. Context discoloration moderate, rose then ochre, stipe base slightly bulbous, basidiospores mostly oblong and cylindrical (Q_m_ > 1.65) ……*Amanita lepiotoides* f. *subcylindrospora* Neville & Poumarat
3. Occurrence in Autumn, on siliceous sandy soil …………………………..*Amanita curtipes* E.-J. Gilbert
3. Occurrence from late Winter (rarely in January) to Spring, also early Summer …………………..4
4. Heavy habit, semi-hypogeous, typically with an earthy odour, associated with *Cistus*, basidiospores mostly ellipsoid to oblong (1.60 < Qm < 1.80) ……..*Amanita ponderosa* Malençon & Heim f. *ponderosa*
4. Medium-sized (pileus diameter generally less than 10 cm) ……………………..5
5. On siliceous sandy soil, associated or not with *Cistus*, pines, etc., basidia with average length < 55 μm ………………………………………………..6
5. On heavy, often naked soil generally of schists, most commonly associated with *Cistus*, basidia with average length > 55 μm ……………..7
6. Relatively small (pileus diameter usually around 5 to 6 cm and below 8 cm), taste of the context indistinct, basidiospores mostly oblong (1.8 < Q_m_ < 2.05) *Amanita curtipes* E.-J. Gilbert
6. Medium-sized (pileus diameter usually between 7 and 10 cm), taste can be slightly of hazelnut, then bitter, basidiospores mostly cylindrical (2.0 < Q_m_ < 2.4)…………. *Amanita pseudovalens* (Neville & Poumarat) Arraiano-Castilho *et al*. var. *pseudovalens* comb. et stat. nov.
7. Odour and/or taste typically distinct, semi-hypogeous habit (*Amanita ponderosa*) ………8
7. Odour indistinct, aftertaste slightly bitter, epigeous habit, pileus margin occasionally with brown scales …….*Amanita pseudovalens* var. *tartessiana* Arraiano-Castilho *et al*. var. nov.
8. Basidiospores broadly oblong (Q_m_ 1.60 – 1.80), strictly thermophile ……*Amanita ponderosa* Malençon & Heim f. *ponderosa*
8. Basidiospores long-ellipsoid (Q_m_ 1.45 – 1.55), also in cooler areas … *Amanita ponderosa* (Gilbert) f. *valens* Neville & Poumarat

### Molecular probes for discrimination from *Amanita ponderosa*

The use of ITS and LSU primers specific for *A. pseudovalens* on the Ode01-Ode16 samples indicated that all DNA extracts from the Luzianes A/B collections (Ode1-8 and Ode10-12) and from the Portalegre collection (Ode16) were *A. pseudovalens* (Fig. 6). All extracts that did not amplify with these primers amplified with the positive control primer sets (not shown). Similar results were obtained with some of the herbarium materials (Fig. S1).

**Fig. 6.**
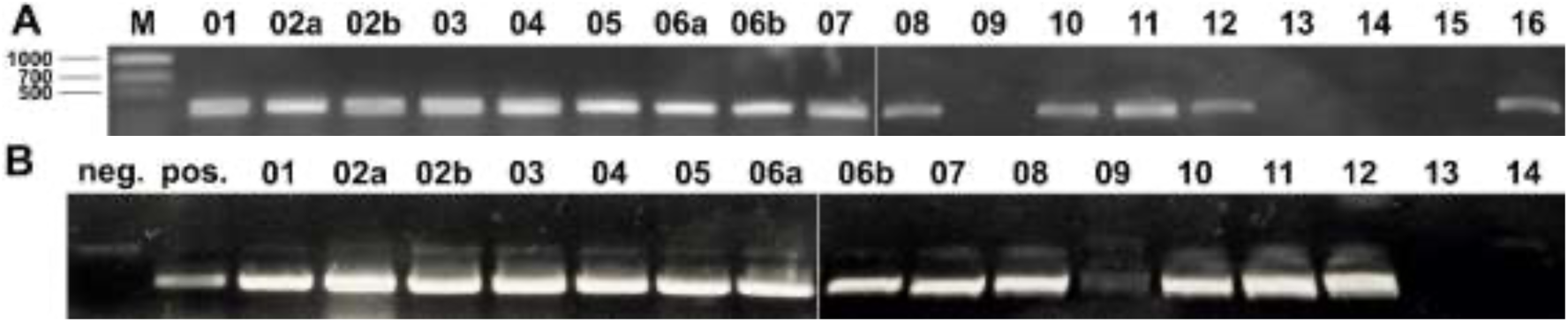
Results of PCR amplifications using discriminant probes for *Amanita pseudovalens*. **A**, ITS region: ITS3–*Aps*Ir3 (all samples positive with concurring control with primers ITS3–ITS4, not shown). **B**, LSU region: NLC2R–*Aps*Lr2 (all positive with control NLC2R–LR5) — Ode15 and Ode16 were negative and positive, respectively, in another amplification (not shown); neg. designates the negative control (water) and pos. the positive control (the P01 extract).

## Discussion

The present study introduces evidence supporting the reclassification of *Amanita curtipes* f. *pseudovalens* to *Amanita pseudovalens* comb. et stat. nov., as well as defining a new variety for the Portuguese collections of the species, *Amanita pseudovalens* var. *tartessiana* var. nov.. The data provide for a clarification of the diagnostic criteria among the European species of *Amidella*, supplemented by the design of a straightforward molecular approach to discriminate the new variety from smaller specimens of *Amanita ponderosa*. The need for this approach is underlined by what appears to be an evolutionary convergence process.

### Taxonomic resolution of the European *Amidella* species

The proposed reclassification to *Amanita pseudovalens* is well supported by the nrDNA sequences, with a net distance of 6.4% to the *A. curtipes* clade (Table 3), ruling against the previous notion that it is merely a robust form of *A. curtipes* (Neville & Poumarat 2009, page 68). This branch within the *Amidella* clade contains the *Amanita pseudovalens* holotype and other herbarium specimens collected in France that conform to its description, as well as an *Amanita ponderosa* lookalike collected in various locations of Southern Portugal, here proposed to belong to *Amanita pseudovalens*. The morphology and ecology of the Portuguese collections is quite distinct from the *A. pseudovalens* original concept (Table 6), such that their conspecificity would not be thinkable without the genetic data. The phylogenetic resemblance suggested by the nrDNA data does not preclude that the new variety is a separate species (Badotti et al. 2017, Vu et al. 2019), but the latter hypothesis will require a definite genetic support.

The literature of *Amidella* in Europe is relatively scarce and hampered by diverging interpretations that remain unresolved to this day, notwithstanding the decisive advances made by Neville & Poumarat (2004). It is clear to us that basidiome size (aside perhaps from the extreme represented by large specimens of *Amanita ponderosa*) is not a pivotal diagnostic character for the group. In this we propose a set of identification keys, where fruiting season, soil texture and associated vegetation are critical for the determination of most European *Amidella* taxa. Thus, it becomes possible to review previous identifications, most notable being those in the extensive study by Castro (1997), compiling collections in Spain and Portugal identified as *Amanita curtipes* (including *A. ponderosa* as a variety of the former, a concept that subsequently has been rejected by Neville & Poumarat (2004), and rebutted by Moreno et al. (2008)). Of the 59 *A. curtipes* collections listed by Castro (1997), at least 24 (in 9 Spanish provinces) occurred between the end of February and mid-July. Although *A. curtipes* is not exclusively autumnal (as exemplified by our specimen M02, see also the proposed key 6), a verification of the *Amanita pseudovalens* identity of some of these collections, reckoned by Neville & Poumarat (2004, page 660), would have a high chorological interest. Probing those collections with our diagnostic primers (Tables 6 and S1) should provide a prompt answer and, together with the field annotations on the soil and vegetation that may exist for them, along with the individual statistics on basidiospore dimensions, could elucidate those collections and, in the ones confirmed to be *Amanita pseudovalens*, enrich the knowledge on this species. Of note, the ‘Q = (1,5)-2-2,2’ reported by Castro (1997) is suggestive of the Q values described for *Amanita pseudovalens* var. *pseudovalens* (Table 6), with the lower range (down to 1.5) possibly indicating the presence of other taxa, including the new variety *Amanita pseudovalens* var. *tartessiana*, especially if the soil and vegetation records are concordant. The latter is so similar to comparatively small specimens of *Amanita ponderosa* (Table 6) that only a few nonmolecular characters are available to set them apart (identification key 7), and those characters might be difficult to ascertain in some collections. Thus, any of the smaller *A. ponderosa* collections in that study might prove to be *A. pseudovalens* var. *tartessiana*.

Given its intermediate spore morphology and the sandy texture of the soil where it was collected, the Ode16 specimen is not unlike the *Amanita pseudovalens* autonym.

Concerning other parts of the *Amidella* phylogeny, a few remarks can be made. First, more collections of *Amanita ponderosa* f. *valens* are needed, and their DNAs studied — for one, Ode14 may already belong here, given our measurements of the basidiospores: length 11.8 μm, width 7.8 μm, Q 1.53 (averages of 14 spores). Second, *Amanita lepiotoides* f. *lepiotoides* is not yet represented in the databases, since all *A. lepiotoides* collections represented in Fig. 2 are identifiable as the f. *subcylindrospora* (Alonso-Díaz & Rigueiro-Rodríguez 2019, Arrillaga & Mayoz 2005, Neville & Poumarat 2004); in this regard, two immature collections from Varese, Italy, dated 1999 (Neville & Poumarat 2004), may be good candidates for sequencing. Finally, one collection from Korea identified as *Amanita curtipes* (Genbank accession numbers KM052544 and KU139473) is rather distant from the European sequences of this species (results not shown) and might be a separate taxon.

### Relevance of the molecular markers

In comparison with other European *Amidella* species, it is generally considered that marketed *Amanita ponderosa* specimens are large and sturdy enough to leave no doubts on their identification (see the first entry in the proposed key 4). Even so, our specimens from Portel, picked by traditionally trained mushroom collectors, included one *Amanita pseudovalens* var. *tartessiana* (P01), and later collections have shown that this co-occurrence with *Amanita ponderosa* is far from negligible (Arraiano-Castilho 2013; Fátima Pinho-Almeida, personal communication). In fact, the few characters available so far for discriminating them (key 7) might not be useful to ascertain some collections, either for being difficult to interpret or for being inconstant.

Thus, assessing the ecological, gastronomical, and economic impacts of such mix-ups is a significant question to be tackled. The genetic markers developed in this study seem to be quite stable across the varied materials examined (Fig. S1) and, combined with a simple DNA extraction method such as the one used in the present work (Stirling 2003), they provide a simple, scalable, and effective means toward monitoring the presence of *Amanita pseudovalens* var. *tartessiana* among the *A. ponderosa* collected for human consumption.

### Evolution and plasticity of the new variety

Table 6 shows how much the new variety, *Amanita pseudovalens* var. *tartessiana*, resembles *Amanita ponderosa* much more than the species autonym, and this intraspecific divergence is found both in the microscopy and in the ecology. A likely evolutionary scenario is that the common ancestor of the *curtipes/pseudovalens* clade (Fig. 2) was adapted to sandy soil and producing (sub)cylindrical spores, a combination of characters retained by var. *pseudovalens* while var. *tartessiana* adapted to schists with *Cistus* covering and producing ellipsoid-oblong spores. Other from its lack of large specimens, the *tartessiana* variety has been converging with *A. ponderosa*, and the plasticity of spore morphology observed in this variety (Fig. 3, Table 5 and statistical analyses) may suggest that such process is still underway. More data on the *curtipes/pseudovalens* clade, including the Korean occurrence mentioned above, might shed some light on this process.

A genome-wide exploration of the divergence between the two varieties of *Amanita pseudovalens* may be a productive model for identifying genotypic markers of the evolutionary divergence process and understanding functions associated with it.

## Conclusion

The genetic comparison between the *Amanita* variety described in this study and the type specimens of three forms established by Neville & Poumarat (2004) firmly suggests the genetic rank for *Amanita pseudovalens* comb. et stat. nov., conspecific with the new collections from Southern Portugal, of a diverging ecology, here described as *Amanita pseudovalens* var. *tartessiana* var nov.. These taxonomic novelties helped clarify the diagnosis of European species within the *Amidella* clade, paving the way for a re-evaluation of vernal collections, henceforth supported by diagnostic molecular probes proposed in this study. Evolutionary convergence may contribute to the identification difficulties in the *Amidella* clade.

## Acknowledgments

Mr. Manuel Nobre Francisco de Vale Fontes guided the authors to the Luzianes locations A and B. The authors dedicate this work to the Iberian mycologists Fátima Pinho-Almeida and Marisa Castro.

