## Supplementary information for "The *Amidella* clade in Europe (*Amanita* Pers., *Basidiomycota*: *Amanitaceae*): clarification of the contentious *Amanita valens* and the importance of taxon-specific PCR primers for identification"

Table S1. List of taxon-specific primers, their matching and positive control universal primers and annealing temperatures ($T_a$), used either for sequencing or as molecular markers.

| Target | Region | ID | Strand | Position* | Sequence | $T_a$(C) | matching | control | Comment |
|---|---|---|---|---|---|---|---|---|---|
| *Amanita curtipes* | ITS | *Acu*If1 | plus | 97 | 5’ AGCGGGCATGTGCACG | 63.5 | ITS2 | ITS1 | |
| | | *Acu*If2 | plus | 533 | 5’ TGCATGAAGAGGAGTTTTTGGAC | 54.3 | ITS4 | ITS3 | |
| | LSU | *Acu*Lr1 | minus | 1410 | 5’ GCCAGCCCATTGAATGATCTG | 55.0 | NLC2R | LR3 | |
| *Amanita lepiotoides* | ITS | *Ale*If1 | plus | 276 | 5’ GTTTTATATGAATGGCTATTG | 51.0 | ITS4 | ITS3 | |
| | | *Ale*Ir1 | minus | 572 | 5’ GTCCAACAAACATCTCCAAC | 56.9 | ITS1F | ITS2 | |
| | LSU | *Ale*Lf1 | plus | 1195 | 5’ GAAGTCAGTAGAGTTGGCTG | 57.5 | LR3 | NLC2R | |
| | | *Ale*Lr1 | minus | 1301 | 5’ CCTTAGCTTTTCTCTAGTG | 57.5 | NLC2R | LR3 | |
| *Amanita ponderosa* | ITS | *Apo*If1 | plus | 58 | 5’ TTAAAACTCTGGCAAAG | 49.4 | ITS2 | ITS1 | |
| | | *Apo*If2 | plus | 499 | 5’ GAGTGTCATTCATATTCTC | 50.3 | ITS4 | ITS3 | |
| | LSU | *Apo*Lr1 | minus | 1399 | 5’ CAAAGCCCAGCCTAGTAG | 56.8 | NLC2R | LR3 | |
| *Amanita pseudovalens* | ITS | *Aps*Ir1 | minus | 339 | 5’ TTGTTCATTAACAATTGTCTTTC | 54.7 | ITS1F | ITS4 | *A. curtipes* produces very faint bands; Ode11 is divergent |
| | | *Aps*If1 | plus | 522 | 5’ AGTCTTCTTCGTGACAAG | 53.9 | ITS4 | ITS3 | var. *pseudovalens* has mismatches |
| | | *Aps*Ir2 | minus | 691 | 5’ CTTATTTCATGGTGTAAAACTTTATC | 55.9 | ITS3 | ITS4 | |
| | | *Aps*Ir3 | minus | 731 | 5’ GACACAAATTCATTTAGAAAG | 51.5 | ITS3 | ITS4 | *A. curtipes* produces very faint bands |
| | LSU | *Aps*Lf1 | plus | 887 | 5’ GCAGCGTCTGGTTGTCC | 60.3 | NLC2 | LR0R | *A. curtipes* also positive (only one mismatch) |

| Target | Region | ID | Strand | Position* | Sequence | $T_a$(C) | matching | control | Comment |
|---|---|---|---|---|---|---|---|---|---|
| | | *Aps*Lr1 | minus | 1301 | 5’ CCTTAGCTTTTTCTACCGGTCAGAA | 62.9 | NLC2R | LR3 | |
| | | *Aps*Lr2 | minus | 1401 | 5’ AATGAATGGCCTGGCGA | 59.1 | NLC2R | LR3 | Some *A.curtipes* specimens display a distinctive set of slower-migrating bands |

* refers to the site number of the 5’ terminal on the alignment used for the phylogenetic reconstruction in Figure 2.

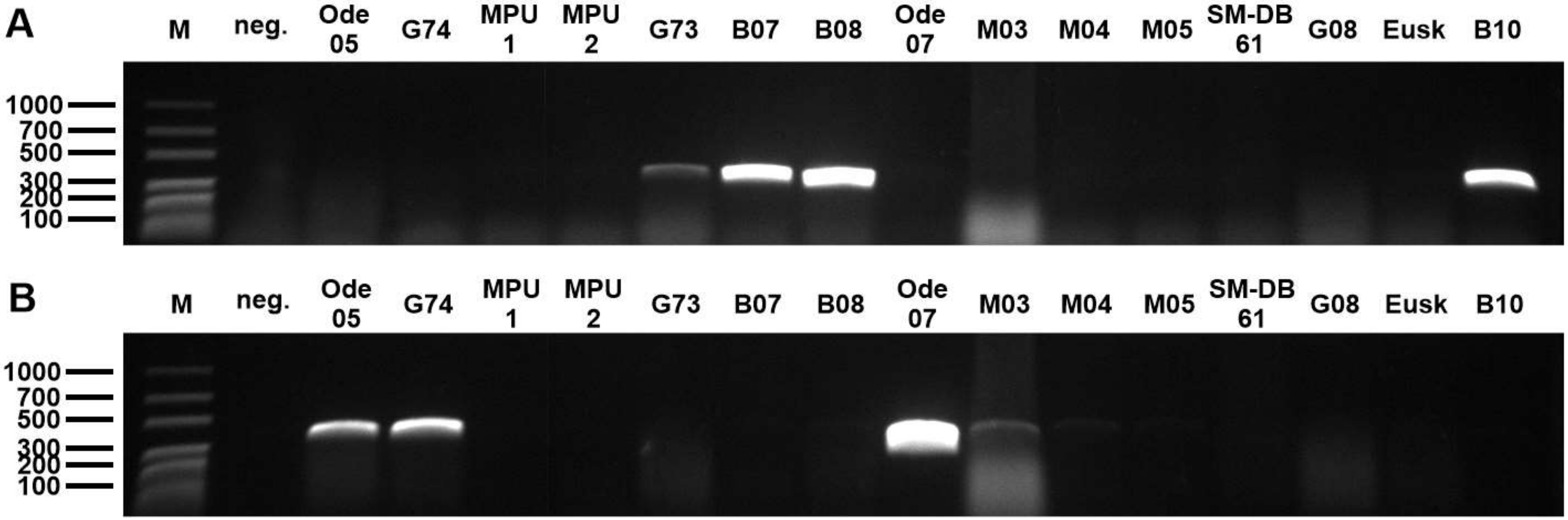


Figure S1. Example of a matched comparison of *Amanita ponderosa* (*Apo*If2-ITS4) and *Amanita pseudovalens* (ITS3-*Aps*Ir3) for the same set of samples representing these two species, along with others of *A. curtipes* and *A. lepiotoides*, and an unknown sample (SM-DB 61). The whole contents of each 20 µL PCR reactions were loaded, to help detect faint signals. Negative controls (neg.) with water instead of DNA. *Amanita ponderosa* represented by the *valens* type (G73) and three samples from Spring 2010 (B07, P08, B10). *Amanita pseudovalens* represented by the *pseudovalens* type (G74), the two MPU samples, Ode05 and Ode07. The *Amanita curtipes* samples (M03, M04 and M05) may produce a weak co-migrating band with the ITS3-ApsIR3 primer pair, and the *A. lepiotoides* samples (G08 and Eusk) were negative in both cases. Positive control reactions were not attempted because herbaria materials may be contaminated with other fungi. M: molecular marker.

Table S2. Usefulness of the taxon-specific primers for the generation of PCR amplicons from herbarium specimens' DNA. ITS-I and ITS-II represent the first and second nrDNA intron, respectively, and LSU represents the proximal region of the 26S nrDNA.

| Taxon | Specimen | ITS-I | ITS-II | LSU | Comments |
|---|---|---|---|---|---|
| *Amanita lepiotoides* f. *subcylindrospora* | G08 | ITS1F-*Ale*Ir1 | *Ale*If1-ITS4B[a] | LR0R-*Ale*Lr1<br>*Ale*Lf1-LR3[a]<br>*Ale*Lf1-LR5 | |
| *Amanita ponderosa* f. *valens* | G73 | *Apo*If1-ITS2_KYO2 | *Apo*If2-ITS4 | NLC2R-*Apo*Lr1 | |
| *Amanita pseudovalens* | G74 | SR6R[a]-*Aps*Ir1 | *Aps*If1-NLC2 | *Aps*Lf1- *Aps*Lr2 | |
| | MPU1 | ITS1F-*Aps*Ir1 | ApsIf1-ITS4 | LR0R-*Aps*LR1<br>NLC2R-*Aps*Lr2 | all regions after reamplification |
| | MPU2 | SR6R-*Aps*Ir1 | *Aps*If1-ITS4 | *Aps*Lf1- *Aps*Lr2 | ITS-II region after reamplification |

[a] https://sites.duke.edu/vilgalyslab/rdna_primers_for_fungi/ (retrieved May 15th 2021)

**Statistical analyses**

<u>Spore measurements</u>

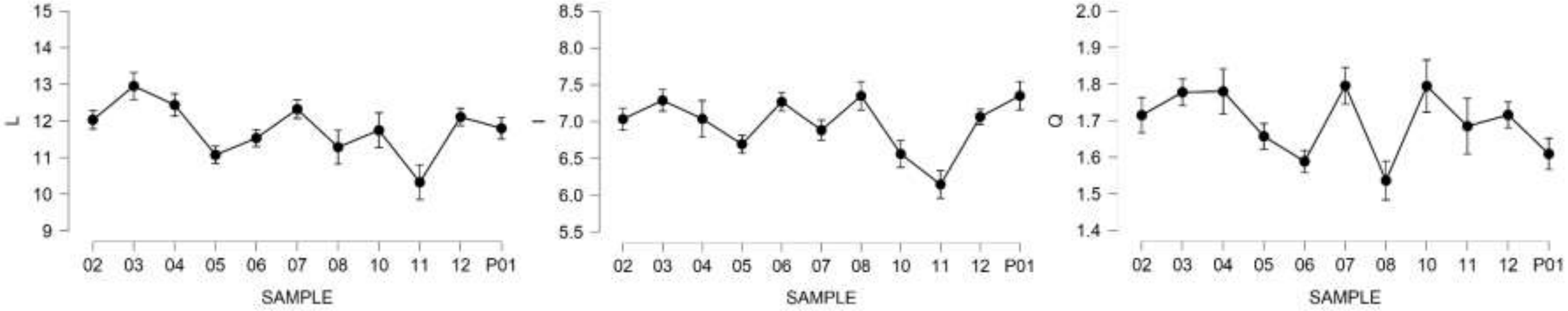


Fig. S2. Descriptives (mean ± SE) plots for basidiospore length (L), width (l) and L/l ratio (Q). Sample identifiers refer to the Ode samples (Spring 2015) and P01 (Spring 2010).

Table S3. Results of the ANOVA tests for basidiospore length (L), width (l) and L/l ratio (Q).

| Measurement | L | l | Q |
|---|---|---|---|
| ANOVA, uncorrected | $F_{10,361}$=19.583*** | $F_{10,361}$=21.031*** | $F_{10,361}$=13.684*** |
| Levene's test (variance homogeneity) | $F_{10,361}$=3.846*** | $F_{10,361}$=3.786*** | $F_{10,361}$=4.314*** |
| ANOVA (Welch correction) | $F_{10,141.108}$=16.760*** | $F_{10,141.158}$=19.401*** | $F_{10,140.459}$=15.368*** |
| Maximum Effect size (Cohen's d) | 2.207*** (Ode03 – Ode11) | 2.463*** (Ode06–Ode11) | 1.907*** (Ode03–Ode08) |

*** P < 0.001

Table S4. Differences between sites for basidiospore length (L), width (l) and L/l ratio (Q).

| Measurement | L | l | Q |
|---|---|---|---|
| Averages ±SE, µm | 12.192±0.085 (A)<br>11.396±0.086 (B) | 7.156±0.040 (A)<br>6.750±0.043 (B) | 1.709±0.012 (A)<br>1.694±0.013 (B) |
| Mean difference (A minus B) ±SE | 0.796±0.121, $t_{340}$=6.573*** | 0.406±0.059, $t_{340}$=6.904*** | 0.015±0.018, $t_{340}$=0.839$^{ns}$ |
| Effect size (Cohen's d) | 0.712 | 0.747 | 0.091 |
| Shapiro-Wilk normality test | W=0.978* (A)<br>W=0.994$^{ns}$ (B) | W=0.986$^{ns}$ (A)<br>W=0.995$^{ns}$ (B) | W=0.987$^{ns}$ (A)<br>W=0.971*** (B) |
| Levene's test (variance homogeneity) | $F_1$=2.313$^{ns}$ | $F_1$=2.894$^{ns}$ | $F_1$=1.029$^{ns}$ |
| Nested ANOVA, between sites | $F_{1,132}$=58.985*** | $F_{1,132}$=64.524*** | $F_{1,132}$=0.937$^{ns}$ |
| Nested ANOVA, within sites | $F_{8,332}$=16.527*** | $F_{8,332}$=16.028*** | $F_{8,332}$=15.068*** |

*** P < 0.001, * P < 0.05, ns P ≥ 0.05

Table S5. Analyses for basidia length.

| Test | Result |
|---|---|
| ANOVA | $F_{7,116}$=2.399, P=0.025 effect size $\omega^2$=0.073 |
| Levene's test (variance homogeneity) | $F_{7,116}$=1.267, P=0.273 |
| Maximum Effect size (Cohen's d) | 1.746, P=0.009 (Ode11–Ode02b) |
| ANOVA without Ode11 | $F_{6,107}$=1.327, P=0.251 effect size $\omega^2$=0.017 |
| Levene's test (variance homogeneity) | $F_{6,107}$=1.256, P=0.284 |

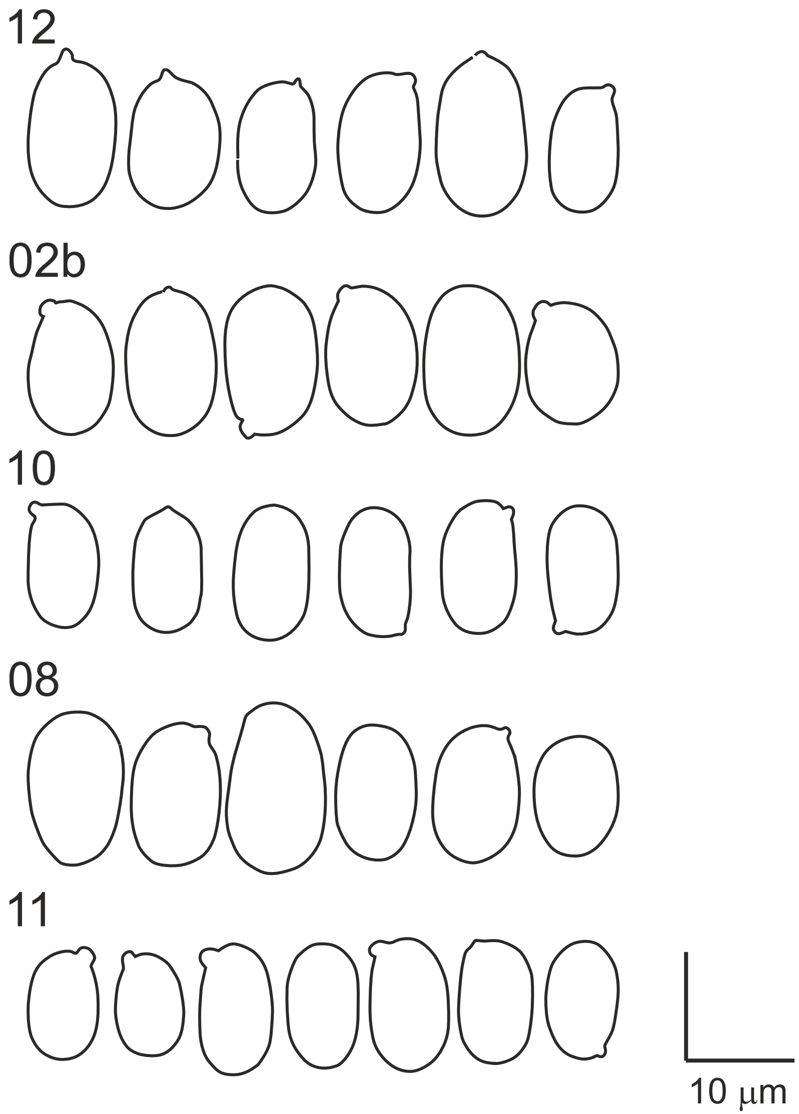

Figure S3. Examples of basidiospore outlines from the Spring 2015 Luzianes collections.
